# Progression of the late-stage divisome is unaffected by the depletion of the cytoplasmic FtsZ pool

**DOI:** 10.1101/2020.06.30.180018

**Authors:** Nadine Silber, Christian Mayer, Cruz L Matos de Opitz, Peter Sass

## Abstract

Cell division is a central and essential process in most bacteria, and also due to its complexity and highly coordinated nature, it has emerged as a promising new antibiotic target pathway in recent years. We have previously shown that ADEP antibiotics preferably induce the degradation of the major cell division protein FtsZ, thereby primarily leading to a depletion of the cytoplasmic FtsZ pool that is needed for treadmilling FtsZ rings. To further investigate the physiological consequences of ADEP treatment, we here studied the effect of ADEP on the different stages of the FtsZ ring in rod-shaped bacteria. Our data reveal the disintegration of early FtsZ rings during ADEP treatment in *Bacillus subtilis*, indicating an essential role of the cytoplasmic FtsZ pool and thus FtsZ ring dynamics during initiation and maturation of the divisome. However, progressed FtsZ rings finalized cytokinesis once the septal peptidoglycan synthase PBP2b, a late stage cell division protein, colocalized at the division site, thus implying that the concentration of the cytoplasmic FtsZ pool and FtsZ ring dynamics are less critical during the late stages of divisome assembly and progression.

## Introduction

Cell division is a vital process in most bacteria and ensures the generation of progeny, usually by yielding equal daughter cells. In rod-shaped bacteria, such as *Bacillus subtilis*, cell division occurs at midcell, driven by FtsZ and orchestrated by a diverse set of proteins together constituting the divisome^1,2^. As the pacemaker of cell division, FtsZ self-polymerizes into protofilaments, which are characterized by additional lateral interactions resulting in higher order polymer assemblies^3^, to finally build the FtsZ ring at the future division site. Here, the FtsZ ring acts as the scaffold for other divisome members. During polymerization, the T7 loop of one FtsZ subunit inserts into the GTP binding site of the next subunit, thereby triggering hydrolysis of GTP to GDP and favoring the disassembly of FtsZ protofilaments^4,5^. Hence, there is a constant and rapid exchange of FtsZ subunits between the dynamic FtsZ ring and the cytoplasmic pool of FtsZ^6-8^. As the cell cycle proceeds, the divisome constricts and synthesizes septal peptidoglycan to allow for septum formation and eventually cytokinesis. Consequently, the process of divisome assembly and progression can be separated into different steps, for example, with regard to the attachment of the FtsZ ring to the membrane, the extent of FtsZ ring constriction, or the consecutive arrival of early and late cell division proteins^7,9,10^. Over the last years, principally two mechanisms have been discussed on how the force required for cytokinesis is generated, either by the chemical energy of FtsZ-dependent GTP hydrolysis^11,12^, or alternatively by peptidoglycan synthesis^11,13^. Recently, FtsZ treadmilling, the GTP-dependent dynamic exchange of FtsZ from the cytoplasmic pool with the FtsZ ring, has been reported to drive divisome progression and constriction in rod-shaped bacteria^14,15^. In cocci such as *Staphylococcus aureus*, however, cell division mainly occurs in two steps in this context, an initial step involving FtsZ treadmilling and a second step that increasingly depends on peptidoglycan synthesis^16^. In this study, we set out to further investigate the role of the cytoplasmic FtsZ pool and FtsZ ring dynamics on the distinct steps of septum formation in rod-shaped bacteria.

## Results and Discussion

To examine the effect of the abundance of FtsZ on divisome assembly and progression, we employed antibiotics of the ADEP class as tools to rapidly modulate the cytoplasmic pool of FtsZ in the model organism *B. subtilis*. ADEP deregulates the bacterial caseinolytic protease, activating its dormant core ClpP for the untimely degradation of FtsZ^17,18^. ADEP incubation thus leads to an impressive filamentation phenotype of *B. subtilis* at concentrations close to the minimal inhibitory concentration (MIC)^18,19^. Very recently, we showed that ADEP-ClpP preferably targets the N terminus of monomeric FtsZ, leading to unfolding and degradation of the FtsZ N-terminal domain^20^. Intriguingly, N-terminal degradation was prevented upon nucleotide binding to FtsZ, most probably due to a stabilization of the FtsZ protein fold. Hence, at ADEP concentrations resulting in a filamentation phenotype, ADEP primarily leads to a depletion of the cytoplasmic pool of nucleotide-free FtsZ in the bacterial cell^20^, thus reducing the FtsZ concentration below the critical level needed for FtsZ ring formation^21,22^ and continuously removing available FtsZ required for FtsZ ring dynamics (Fig. 1a). In line with this, a recent model on cell-size homeostasis in rod-shaped bacteria described a direct correlation of FtsZ expression and accumulation on septal constriction initiation^23^. Therefore, ADEP was considered instrumental to investigate the role of the cytoplasmic FtsZ pool and FtsZ ring dynamics during divisome formation and progression. To do so, we first tested the effect of ADEP on polymerized FtsZ (in the presence of GTP) (Fig. 1b). Once assembled into protofilaments, FtsZ substantially resisted the degradation by ADEP-ClpP. It may thus be hypothesized that, if divisome assembly and constriction fully depend on the cytoplasmic FtsZ pool and FtsZ ring dynamics, ADEP treatment should result in the disintegration of early as well as late stage divisomes, or at least, the progression of late stage divisomes should be halted. To test this hypothesis, we conducted time-lapse and super-resolution fluorescence microscopy experiments with ADEP-treated *B. subtilis* strain 2020 using filamentation concentrations of the antibiotic. This strain additionally expresses FtsZ fused to GFP from an ectopic locus. For microscopy, ADEP-treated bacteria were mounted and then further grown on microscopy slides coated with ADEP-supplemented agarose (approximate division time of bacteria on microscopy slides was 30-45 minutes). Of note, ADEP leads to a depletion of the FtsZ pool within 15-20 minutes, i.e. the cellular concentration of FtsZ is reduced by more than half within 5 minutes of ADEP treatment, whereas biomass increase and metabolism in general remain unaltered^18,20^. Therefore, the elapsed time between the addition of ADEP to the cells and initial image acquisition was set to 15-20 minutes, ensuring that image acquisition started under FtsZ-depleted conditions. By monitoring FtsZ ring formation over time, we observed that ADEP inhibited the initiation of FtsZ ring assembly, and early FtsZ rings that had just been formed disintegrated. Here, the presence of early FtsZ rings at the beginning of the experiment and their soon disintegration indicate that imaging started just before ADEP effects on FtsZ rings could visually be observed. However, in contrast to early FtsZ rings, more progressed FtsZ rings constricted and finished septum formation, apparently being less sensitive to ADEP treatment, finally yielding two separated daughter cells (Fig. 2ab, Supplementary Fig. 1 and 2, Supplementary Movies 1-3). Following up on this, we investigated whether the arrival of the late stage cell division protein PBP2b^9^, a septal peptidoglycan synthase, would coincide with a successful constriction of progressed FtsZ rings during ADEP-treatment. By using *B. subtilis* strain CM03, which allows for the concomitant expression of mCherry-FtsZ and GFP-PBP2b, we observed that early FtsZ rings disintegrated prior to the arrival of PBP2b. On the contrary, the divisome consistently finalized cell division after PBP2b had substantially arrived at the septum area (Fig. 3ab, Supplementary Fig. 3, Supplementary Movie 4). It may therefore be hypothesized that the arrival or activity of late stage cell division proteins, such as peptidoglycan synthases, may substantially support or even trigger divisome progression in rod-shaped bacteria similar as previously suggested for coccoid bacteria, such as *S. aureus*^16^.

**Fig. 1:**
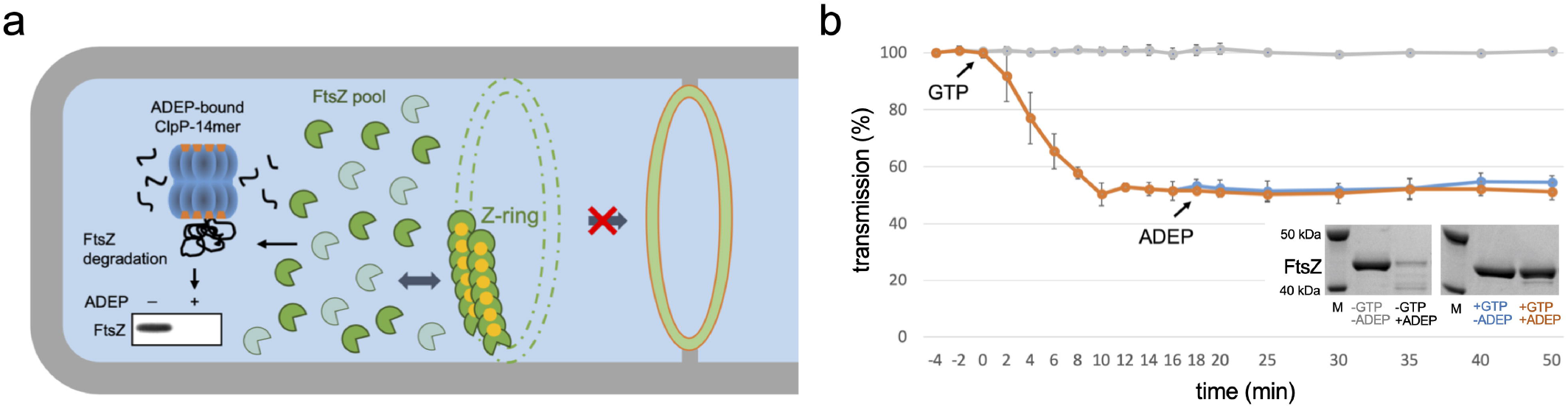
ADEP antibiotics lead to a reduction of the cytoplasmic FtsZ pool. **a**, Schematic of ADEP-dependent degradation of FtsZ. ADEP (orange) activates bacterial ClpP peptidase (blue) for untimely protein degradation, and nucleotide-free, monomeric FtsZ (dark green) represents a preferred protein substrate for ADEP-ClpP^18,20^. As a consequence, the cytoplasmic FtsZ pool is depleted (indicated by the shift from dark to light green), whereas GTP-bound FtsZ (GTP in yellow) is stabilized against proteolytic attack at antibiotic concentrations close to the MIC. FtsZ ring assembly relies on the dynamic exchange of FtsZ subunits between the FtsZ ring and the cytoplasmic FtsZ pool (indicated by dark blue arrow). In the presence of ADEP, the depletion of the FtsZ pool ultimately results in an inhibition of FtsZ ring formation and cell division, finally leading to bacterial cell death. However, it remained unresolved, whether only early or also later stages of the FtsZ-ring/divisome are affected by ADEP treatment. **b**, *In vitro* FtsZ polymerization and FtsZ degradation by ADEP-activated ClpP. Light transmission analyses of GTP-dependent FtsZ polymerization *in vitro* followed by incubation with a ClpP reaction mixture in the absence or presence of ADEP and/or GTP (-GTP/-ADEP, in grey; +GTP/-ADEP, in blue; +GTP/+ADEP, in orange). Here, polymerized FtsZ is not affected by incubation with ADEP-ClpP. The graphs show mean values of two independent experiments, bottom and top values of indicator bars show the individual data points of each respective replicate. Source data underlying the graphs is presented in Supplementary Table 1. As an independent control, we determined FtsZ protein amounts of samples that were incubated with or without GTP for 120 min in the absence or presence of ADEP. DMSO was used as an untreated control in all assays. Representative SDS-PAGE images of triplicates are depicted. Source data of the full, uncropped gel image is presented in Supplementary Fig. 4.

**Fig. 2:**
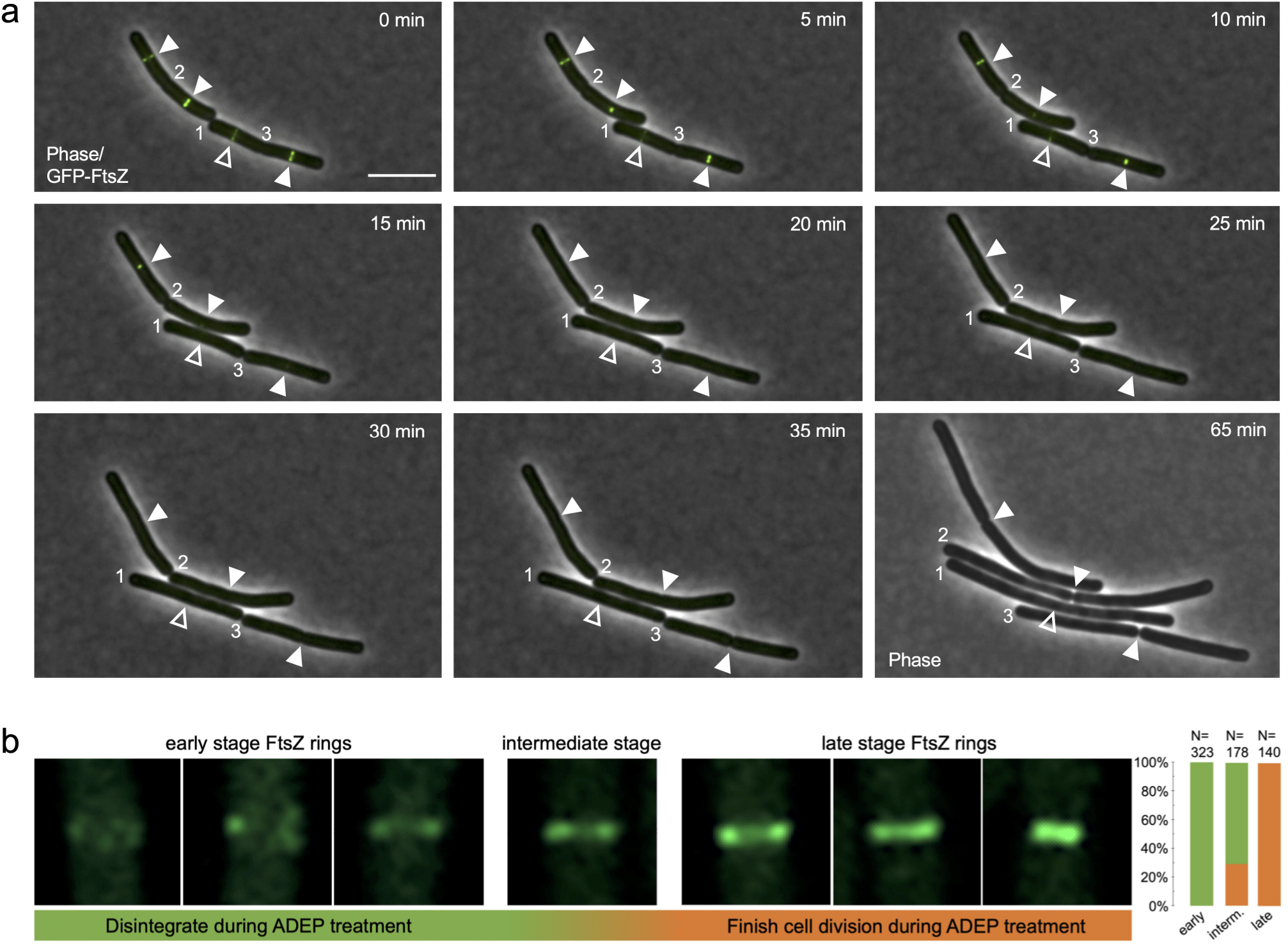
Depletion of the FtsZ pool differently affects distinct stages of the FtsZ ring. **a**, Time-lapse fluorescence microscopy of exponentially growing *B. subtilis* 2020 cells treated with 0.25 µg/ml ADEP2. Overlaid fluorescence and phase contrast images show the localization of GFP-tagged FtsZ (in green) and the progression of FtsZ rings over time. The micrographs indicate that early FtsZ rings (open triangles) disintegrate during ADEP treatment while more progressed FtsZ rings (closed triangles) constrict and finish septum formation to yield two separated daughter cells. Numbers indicate previously finished septa. For clarity, numbers remain positioned to the corresponding cell pole of the daughter cell on the right. A phase contrast image of bacterial cells after 65 min is included at the end of the series to prove failure or success of septum formation. Overlay (0-35 min) or single channel (65 min) images are provided. Scale bar, 5 µm. Images are representative of at least three biological replicate cultures of *B. subtilis* 2020 with >600 FtsZ rings analyzed over time. A time-lapse video is provided by Supplementary Movie 1. **b**, Super-resolution fluorescence microscopy of different stages of FtsZ ring formation in *B. subtilis* 2020. While all of the early stage FtsZ rings disintegrate upon ADEP treatment (100%, N=323), all late stage FtsZ rings further constrict and finalize septum formation (100%, N=140). Intermediate stage FtsZ rings, which were in transition from early to late stages, show a heterogeneous behavior with 71% of FtsZ rings abrogating division and 29% further constricting and finalizing septum formation (N=178). Of note, we deliberately analyzed immediate daughter cells comprising both early and further progressed FtsZ rings, of which all late stage FtsZ rings finalized division while all early FtsZ rings did not, thus making it unlikely that the cells with late stage FtsZ rings may by chance happened to have considerably higher FtsZ levels compared to cells with early FtsZ rings. This is further supported by the fact that ADEP leads to a rapid depletion of the FtsZ pool within 15-20 minutes, where potential minor differences of the FtsZ level may be negligible in relation to the doubling time of the bacteria of approximately 30-45 minutes on microscopy slides. Images are representative of at least three biological replicate cultures. Source data underlying the graphs is presented in Supplementary Fig. 5.

**Fig. 3:**
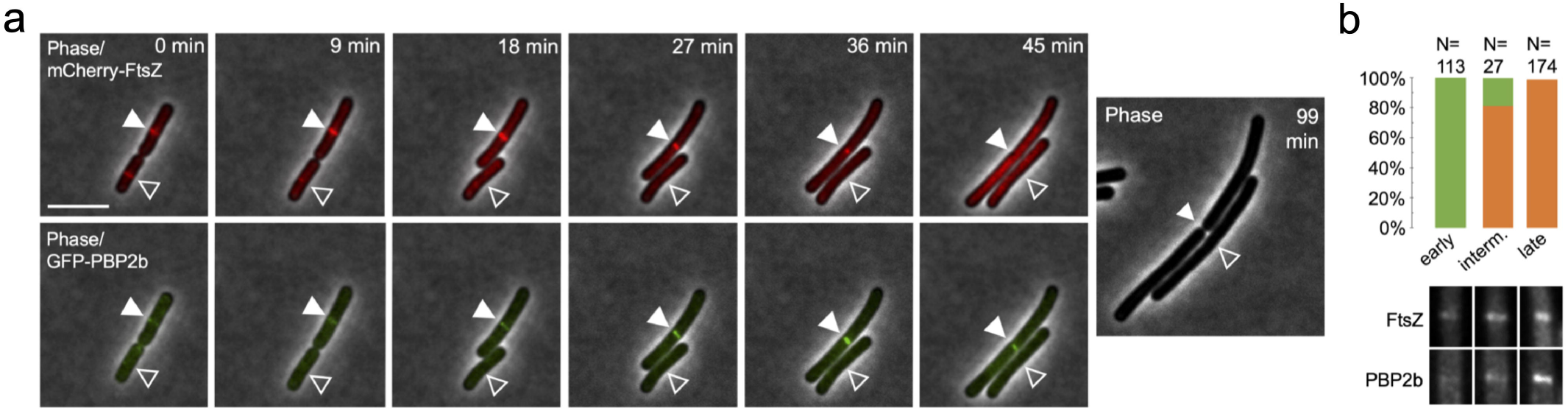
The late-stage divisome progresses independent of the depletion of the FtsZ pool. **a**, Time-lapse fluorescence microscopy of exponentially growing *B. subtilis* CM03 cells treated with 0.125 µg/ml ADEP2. Overlaid fluorescence and phase contrast images show the localization of mCherry-FtsZ or GFP-PBP2b, as indicated, during ADEP treatment over time. The micrographs show that the divisome consistently succeeds to finalize cell division (closed triangles) in the presence of ADEP in situations when GFP-PBP2b is substantially detected at the septum area. In contrast, early stage FtsZ rings, prior to the visible arrival of GFP-PBP2b at the septum, disintegrate during ADEP treatment (open triangles). For clarity, a phase contrast image of bacterial cells after 99 min is included at the end of the series to prove failure or success of septum formation. Scale bar, 5 µm. Images are representative of at least three biological replicate cultures of *B. subtilis* CM03 with >300 septa/FtsZ rings monitored over time. A time-lapse video is provided by Supplementary Movie 4. **b**, Quantification analysis assessing divisome progression and success of cell division in cells with or without clear foci of PBP2b. In the absence of a substantial signal of GFP-PBP2b at the divisome, early stage FtsZ rings disintegrate and cells fail to divide upon ADEP treatment (100%, N=113), however, after substantial localization of GFP-PBP2b, the divisome progresses and cells finalize septum formation and division (100%, N=174). In the transition phase of initial GFP-PBP2b localization, cells show a heterogeneous behavior with 18.5% of the divisomes abrogating division and 81.5% further constricting and finalizing division (N=27). Bottom panels show representative fluorescence microscopy images of the different stages corresponding to the graphs in the upper panel. Images are representative of at least three biological replicate cultures of *B. subtilis* CM03. Source data underlying the graphs is presented in Supplementary Fig. 6.

At the same time, Whitley and colleagues published a preprint reporting a related observation regarding the role of FtsZ treadmilling during cell division, however, using the antibiotic PC190723. In contrast to ADEP, PC190723 stabilizes FtsZ bundles and halts FtsZ dynamics instead of depleting FtsZ by fast degradation as occurs in ADEP-treated cells. In line with our observations, FtsZ treadmilling had a dispensable function in accelerating septal constriction rate in their study, while it was critical for assembling and initiating the bacterial divisome^24,25^.

In conclusion, our data imply distinct stages during FtsZ ring initiation, maturation and constriction in rod-shaped bacteria with regard to the role of the cytoplasmic FtsZ pool as well as the sensitivity to ADEP antibiotics. Clearly, during ADEP treatment it is distinguished between early and more progressed divisomes, thereby adding another level of complexity to the elaborate mechanism of ADEP action. Also, our data support a two-step model of cell division in *B. subtilis* (Fig. 4), in which the late-stage divisome is substantially less sensitive to a depletion of the cytoplasmic FtsZ pool, and thus less dependent on FtsZ ring dynamics, in contrast to initial assembly and early stage FtsZ rings.

**Fig. 4:**
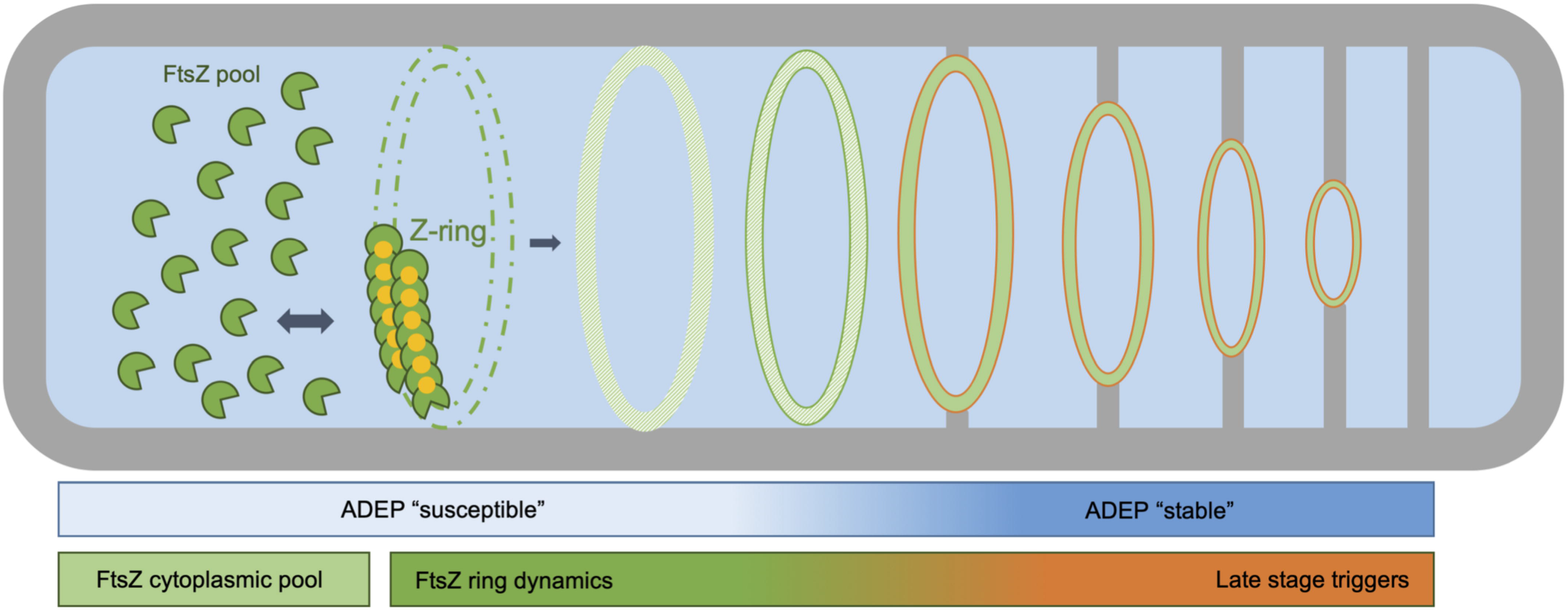
Two-step model of FtsZ ring assembly and progression. At antibiotic concentrations close to the MIC, ADEP treatment essentially depletes the cytoplasmic pool of FtsZ^20^, thereby continuously removing available FtsZ that is needed for FtsZ ring formation and dynamics^14^. Our data reveal that ADEP treatment leads to the disintegration of early FtsZ rings (framed green) in *B. subtilis*, while more progressed FtsZ rings (framed red) resist degradation and finish septum formation as well as daughter cell separation, suggesting distinct stages during FtsZ ring initiation and progression. Therefore, our results support a two-step model of FtsZ ring and divisome progression in *B. subtilis*. In the initial stage, FtsZ ring formation essentially relies on the cytoplasmic FtsZ pool and FtsZ ring dynamics, indicated by the disintegration of early FtsZ rings upon ADEP-dependent depletion of the cytoplasmic FtsZ pool. In the later stage, the more progressed divisome is considerably less sensitive to a depletion of the cytoplasmic FtsZ pool, implying that other triggers take over to drive divisome progression, for example, peptidoglycan synthases that arrive at the divisome during the later stages of cell division.

## Methods

### Protein purification

FtsZ and ClpP proteins were derived from *B. subtilis* 168 (*trpC2*; wild-type strain; NC_000964.3) and were expressed as C-terminally His6-tagged proteins in *E. coli* BL21(DE3) harboring the respective expression plasmid^18,20^. Expression cultures were grown in lysogeny broth (LB) containing ampicillin (100 µg/ml) to an optical density at 600 nm (OD600) of 0.6. Then protein expression was induced by the addition of isopropyl-β -D-thiogalactopyranoside (IPTG) to a final concentration of 1 mM. Expression cultures were harvested after 4 hours followed by standard protein purification steps under native conditions using Ni-NTA affinity chromatography. Quantity and quality of the purified proteins were verified by Bradford assay (using bovine serum albumin as control), Nanodrop spectrophotometry (Nanodrop Technologies) and SDS-PAGE. We have shown previously that purification-tags have no effect on FtsZ degradation or enzyme activity of both proteins^20^.

### *In vitro* FtsZ polymerization and degradation assays

For GTP-dependent polymerization of FtsZ *in vitro*^26,27^, purified *B. subtilis* 168 FtsZ (25 µM) was preincubated in polymerization buffer (50 mM MES/NaOH, pH 6.5, 50 mM KCl, 10 mM MgCl_2_) for 10 min on ice. After initial incubation for 4 min at 37 °C allowing baseline correction, GTP was added to the respective reaction mixture to a final concentration of 1 mM. Then, reaction mixtures were transferred into a photometer cuvette for monitoring light transmission at 400 nm and 37 °C over time. For *in vitro* degradation of polymerized FtsZ, 12 µM ADEP2^17,28^ (or equal volume of DMSO as a control) and 1.5 µM purified ClpP protein (monomer concentration) were added to the polymerization reaction after 16 min, as indicated, and light transmission was further monitored for 34 min. As an independent control, FtsZ (4 µM) was incubated with 1 mM GTP in activity buffer (50 mM Tris/HCl pH 8, 25 mM MgCl2, 100 mM KCl, 2 mM DTT) at 37 °C for 30 min. Then 1.5 µM ClpP (monomer concentration) and 3.75 µM ADEP (or equal volume of DMSO as a control) were added to the reaction mixtures that were further incubated at 37 °C. Samples were taken after 120 min and were analyzed via SDS-PAGE using standard techniques^18,20^.

### Cloning strategy

For colocalization studies of FtsZ with PBP2b (mCherry-FtsZ/GFP-PBP2b), strain *B. subtilis* CM03 was constructed as follows. Plasmid pHJCM03 was generated using the coding sequence of *pbpB* that was amplified from chromosomal DNA of *B. subtilis* 168 (*trpC2*; wild-type strain; NC_000964.3) *via* PCR using the following primers: oCM07 5’-GGAAGCGGCTCAGGCTCCCGGATCCACATTCAAATGCCAAAAAAGAATAAATTTATG-3’, oCM08 5’-CGCGGCCGCTCTAGAACTAGAATTCTTAATCAGGATTTTTAAACTTAACCTTG-3’. The amplicon was then ligated into the linearized plasmid pHJCM01^19^ by a Gibson isothermal reaction. The resulting plasmid was transformed into *B. subtilis* CM01 (*trpC2 cat aprE::Pspac-mcherry-ftsZ*)^19^ to give strain CM03 (*trpC2 cat aprE::Pspac-mcherry-ftsZ; spc amyE::Pxyl-msfgfp-pbpB*).

### Super-resolution and time-lapse fluorescence microscopy

Cells of *B. subtilis* 2020 (*trpC2 spc amyE::Pxyl-gfp-ftsZ*)^29^ or *B. subtilis* CM03 (*trpC2 cat aprE::Pspac-mcherry-ftsZ; spc amyE::Pxyl-msfgfp-pbpB*) were grown at 37 °C to early-exponential phase (OD600 of 0.1). For the expression of GFP and mCherry fusion proteins, *Pxyl* and *Pspac* promoters were induced using 0.1-0.2% xylose and 0.1 mM IPTG, respectively. Then, cells were pre-incubated in lysogeny broth (LB) supplemented with 0.125-0.25 µg/ml ADEP2 for 10 minutes or DMSO as a control. Bacteria were transferred onto microscopy slides using gene frames (Life Technologies) and a thin film of 1.5% agarose in 25% LB containing 0.125-0.25 µg/ml ADEP2 as well as according concentrations of the inducer compounds xylose and/or IPTG. Doubling times of bacteria on microscopy slides were in the range of 30-45 minutes. Phase contrast and fluorescence images were taken at distinct time points as indicated. Elapsed time between sampling and image acquisition was approximately 5-10 minutes. Of note, the ADEP2 concentration used here results in filamentation of *B. subtilis*, resembling a phenotype that is due to fast FtsZ degradation, i.e. the cellular amount of FtsZ is reduced by more than half within 5 minutes once ADEP-ClpP is fully engaged, whereas biomass increase and metabolism in general remain unaltered^18^. Since the addition of ADEP leads to an overall depletion of the FtsZ pool within 15-20 minutes^18,20^, image acquisition was started at the assumed onset of detectable FtsZ pool reduction (correlating to approximately half the bacterial doubling time on microscopy slides), which is further indicated by the presence of early FtsZ rings at the start of imaging as well as their soon disintegration. For image analyses, we deliberately monitored bacterial cells or small cell clusters (i.e. immediate daughter cells) that showed both early as well as late FtsZ rings, thereby ensuring similar conditions of ADEP action on the distinct stages of the FtsZ ring and the divisome in the analyzed bacterial cells. Super-resolution images were recorded using a Zeiss Axio Observer Z1 LSM800 equipped with an Airyscan detector and a C Plan-Apo 63x/1.4 Oil DIC objective (Zeiss, Germany). Images were processed using the ZEN2.3 image analysis software package (Zeiss). Time-lapse micrographs were obtained using a Nikon Eclipse Ti automated microscope equipped with a Perfect Focus system (Nikon Instruments Europe BV, Netherlands), an Orca Flash 4.0 camera (Hamamatsu, Photonics, Japan) and CFI Plan-Apo DM 100x/1.45 Oil Ph3 objective (Nikon). Image acquisition and analysis were performed via the NIS elements AR software package (Nikon).

### Statistics and Reproducibility

Microscopy and *in vitro* FtsZ degradation results were each obtained from at least three independent experiments and biological samples. The results describing the effect of *in vitro* degradation of FtsZ polymers on light transmission (Fig. 1b) are based on two independent experiments. For experiments using samples of small sizes (Fig. 1b, N=2) mean values were calculated and depicted, indicating individual data points via the bottom and top values of the respective indicator bars. For the evaluation of the fate of divisomes (localization of FtsZ and PBP2b) upon ADEP treatment, large numbers of divisomes (>950) from a total of twelve independent biological samples were examined. Total values and percentages are described (Supplementary Fig. 5 and 6).

## Data availability

Additional datasets generated and/or analyzed during the current study are available from the corresponding author on reasonable request.

## Acknowledgments

We thank Heike Brötz-Oesterhelt for providing ADEP2, as well as Leendert Hamoen, Jeff Errington and Kürşad Turgay for the kind gift of strains and plasmids. We also thank Anne Berscheid for critically reading and discussing the manuscript.

The authors appreciate funding by the Deutsche Forschungsgemeinschaft (DFG, German Research Foundation): Project-ID 398967434 (TRR 261-A02), the German Center for Infection Research (DZIF; TTU 09.815), as well as support by infrastructural funding from the Cluster of Excellence EXC 2124 Controlling Microbes to Fight Infections.

## Author contributions

N Silber, C Mayer, CL Matos de Opitz and P Sass performed and designed experiments, analyzed data and prepared figures. P Sass conceived and supervised the study, wrote the manuscript and raised funding. All authors discussed data and edited the manuscript.

