## Supplementary Information for "Progression of the late-stage divisome is unaffected by the depletion of the cytoplasmic FtsZ pool"

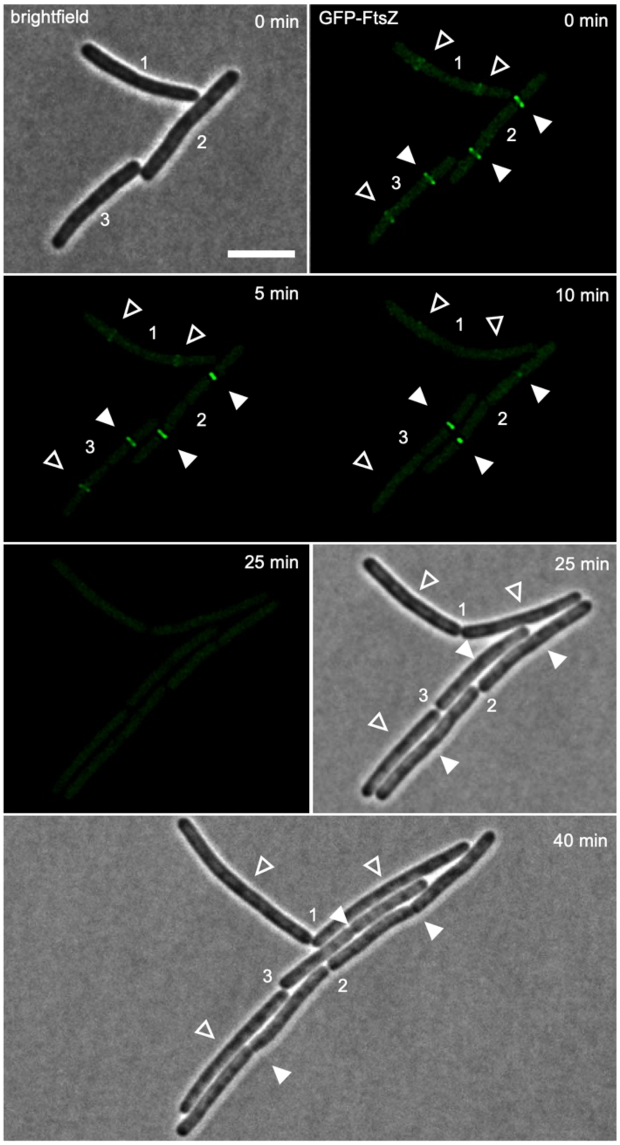

**Supplementary Fig. 1:**

**ADEP-treatment leads to the disintegration of early, but not late stage FtsZ rings.**

Super-resolution fluorescence microscopy of exponentially growing *B. subtilis* 2020 cells treated with 0.25 µg/ml ADEP2. Fluorescence images show the localization of GFP-tagged FtsZ and the progression of FtsZ rings (in green) over time. During ADEP treatment, early FtsZ rings disintegrate (open triangles), while mature FtsZ rings finish septum formation (closed triangles). Numbers indicate already finished, but undivided septa. Scale bar, 5 µm. Images are representative of at least three biological replicate cultures.

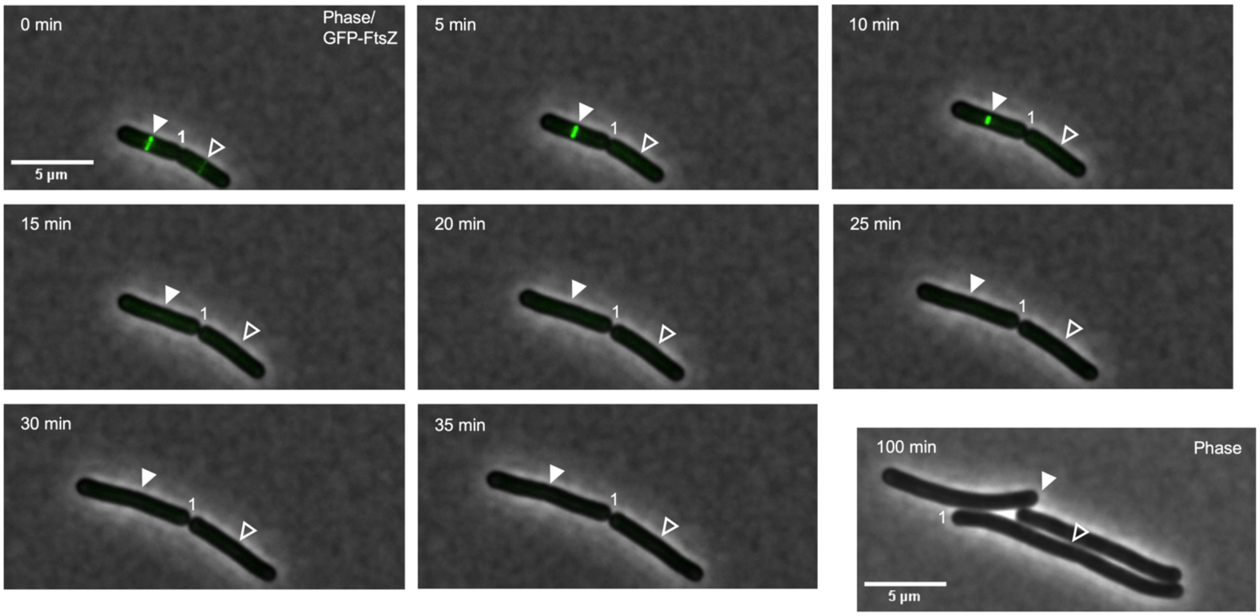

**Supplementary Fig. 2:**

**Effect of ADEP treatment on FtsZ ring formation in *B. subtilis* cells visualized by time-lapse fluorescence microscopy.**

Exponentially growing *B. subtilis* 2020 cells were treated with 0.25 µg/ml ADEP2 and FtsZ ring formation was followed over time. Micrographs show overlays of phase contrast images (greyscale) and GFP fluorescence images (green) indicating FtsZ ring formation at mid-cell. ADEP-treated cells show disintegration of early FtsZ rings (open triangles) while progressed FtsZ rings constrict and finalize septum formation to yield two separated daughter cells (closed triangles). Numbers indicate already finished, but undivided septa. For clarity, numbers remain positioned to the corresponding cell pole of the daughter cell on the right. An additional phase contrast image was acquired after prolonged incubation with ADEP (last image in the series) indicating failure or success of septum formation. Scale bars, 5 µm. Images are representative of at least three biological replicate cultures of *B. subtilis* with >600 FtsZ rings analyzed over time.

A time-lapse video of the image sequence is provided by Supplementary Movie 2.

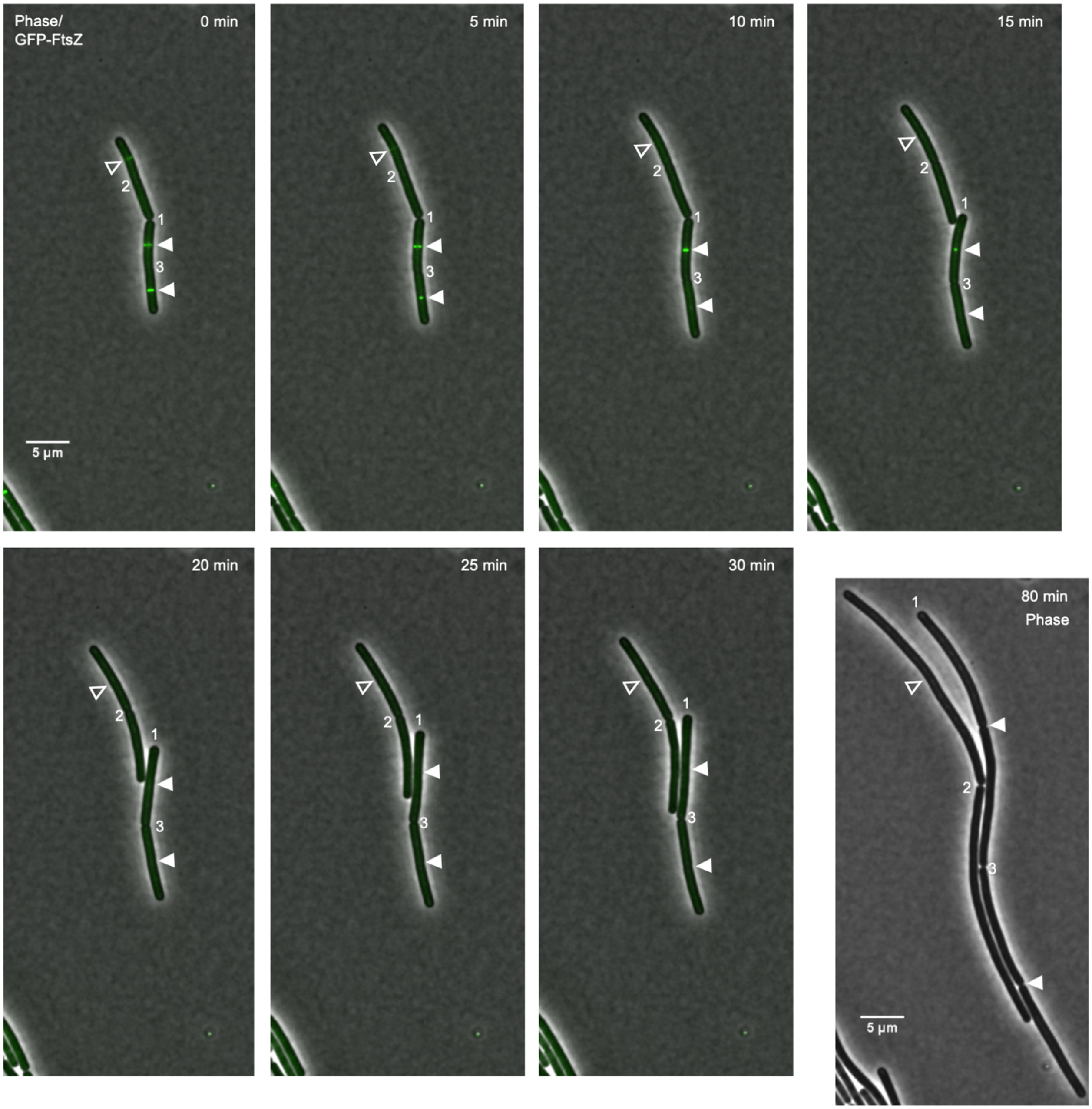

**Supplementary Fig. 2 continued:**

**Effect of ADEP treatment on FtsZ ring formation in *B. subtilis* cells visualized by time-lapse fluorescence microscopy.**

A time-lapse video of the image sequence is provided by Supplementary Movie 3.

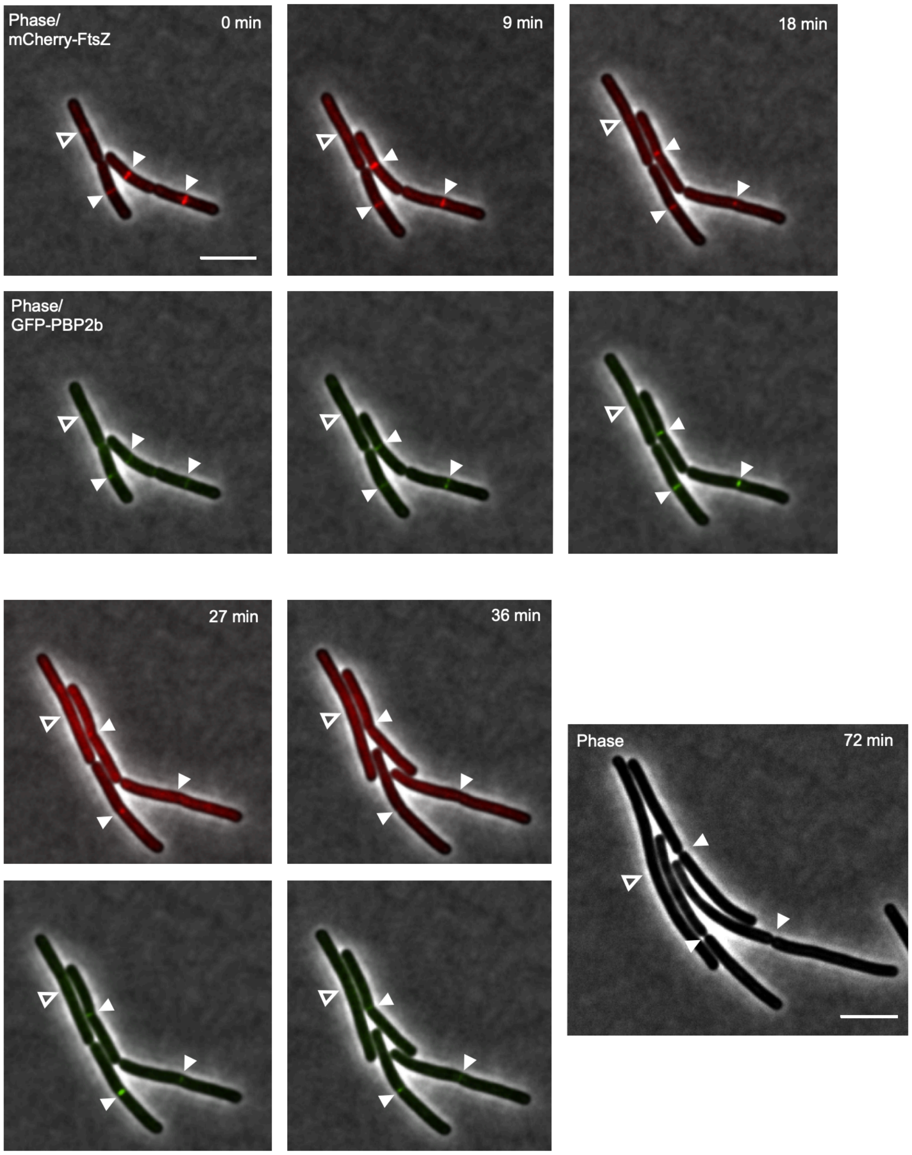

**Supplementary Fig. 3:**

**Colocalization of mCherry-FtsZ and GFP-PBP2b during ADEP treatment of *B. subtilis* cells visualized by time-lapse fluorescence microscopy.**

Overlaid fluorescence and phase contrast images are shown. During ADEP treatment (0.125 µg/ml ADEP2), early FtsZ rings disintegrate (open triangles), while mature FtsZ rings finish septum formation once PBP2b has substantially arrived at the septum (closed triangles). A phase contrast image of bacterial cells after 72 min is included at the end of the series to prove failure or success of septum formation. Scale bars, 5 µm. Images are representative of at least three biological replicate cultures of *B. subtilis* CM03 with >300 FtsZ rings analyzed over time.

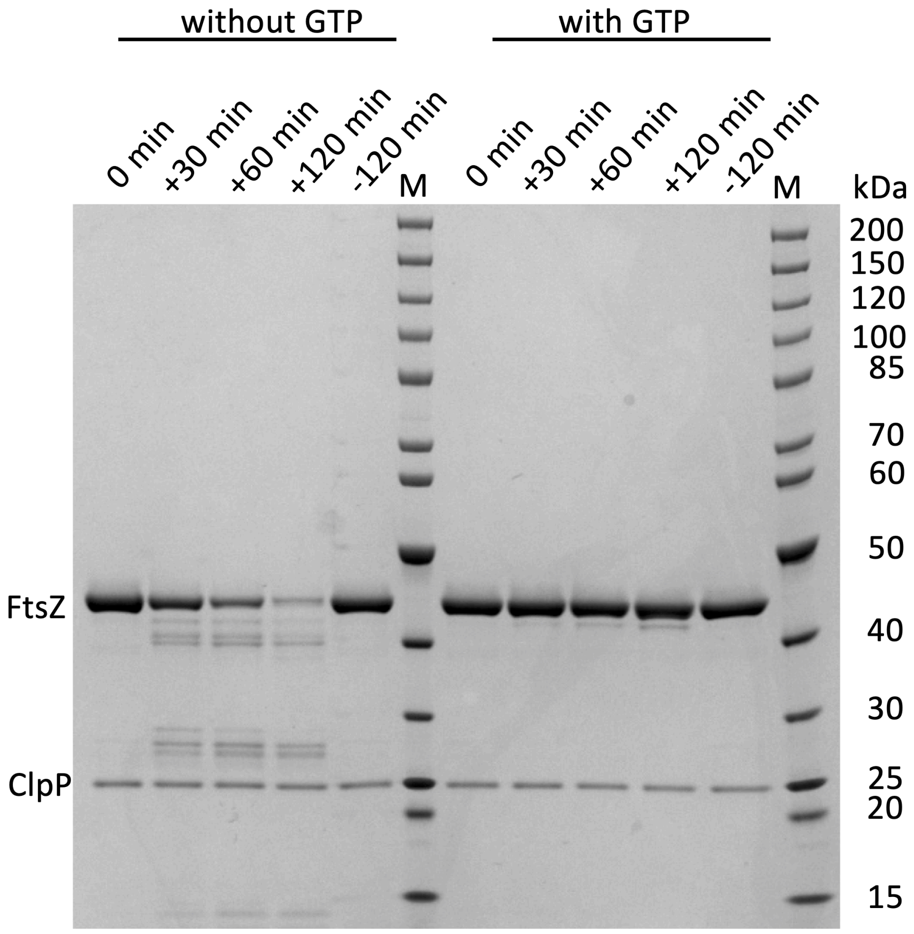

**Supplementary Fig. 4:**

Source data of the full, uncropped gel image according to Fig. 1b.

-, no ADEP; +, plus ADEP; M, molecular marker.

| | early Z-rings | | intermediate Z-rings | | late Z-rings | |
|---|---|---|---|---|---|---|
| | disintegration | constriction | disintegration | constriction | disintegration | constriction |
| experiment 1 | 87 | 0 | 33 | 12 | 0 | 51 |
| experiment 2 | 39 | 0 | 22 | 10 | 0 | 10 |
| experiment 3 | 123 | 0 | 57 | 15 | 0 | 58 |
| experiment 4 | 74 | 0 | 14 | 15 | 0 | 21 |
| sum | 323 | 0 | 126 | 52 | 0 | 140 |
| **sum (total)** | **323** | | **178** | | **140** | |
| [%] | 100 | 0 | 71 | 29 | 0 | 100 |

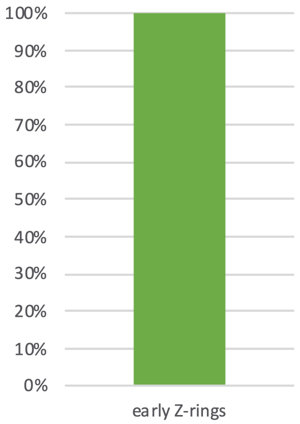

N=323

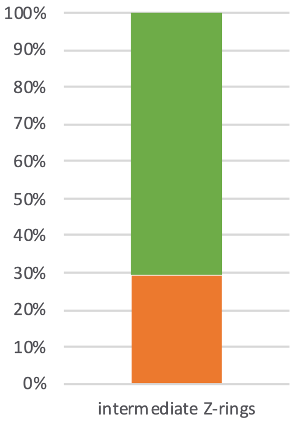

N=178

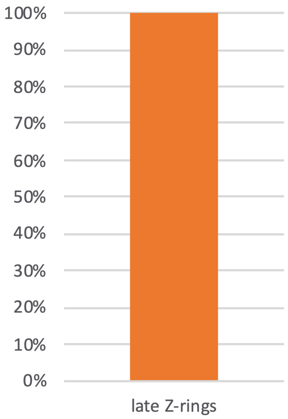

N=140

**Supplementary Fig. 5:**

Source data underlying the graphs in Fig. 2b showing the distribution of either success or failure of FtsZ ring/divisome progression among the cells counted.

| | early Z-rings (no/weak PBP2b) | |
|---|---|---|
| | disintegration | constriction |
| experiment 1 | 4 | 0 |
| experiment 2 | 39 | 0 |
| experiment 3 | 11 | 0 |
| experiment 4 | 2 | 0 |
| experiment 5 | 0 | 0 |
| experiment 6 | 1 | 0 |
| experiment 7 | 8 | 0 |
| experiment 8 | 48 | 0 |
| sum | 113 | 0 |
| **sum (total)** | **113** | |
| [%] | 100 | 0 |

| | intermediate Z-rings (weak, but clear PBP2b) | |
|---|---|---|
| | disintegration | constriction |
| experiment 1 | 1 | 7 |
| experiment 2 | 3 | 13 |
| experiment 3 | 0 | 2 |
| experiment 4 | 0 | 0 |
| experiment 5 | 1 | 0 |
| experiment 6 | n.e. | n.e. |
| experiment 7 | n.e. | n.e. |
| experiment 8 | n.e. | n.e. |
| sum | 5 | 22 |
| **sum (total)** | **27** | |
| [%] | 18.5 | 81.5 |

| | late Z-rings (clear PBP2b signal) | |
|---|---|---|
| | disintegration | constriction |
| experiment 1 | 0 | 65 |
| experiment 2 | 0 | 42 |
| experiment 3 | 0 | 11 |
| experiment 4 | 0 | 44 |
| experiment 5 | 0 | 12 |
| experiment 6 | n.e. | n.e. |
| experiment 7 | n.e. | n.e. |
| experiment 8 | n.e. | n.e. |
| sum | 0 | 174 |
| **sum (total)** | **174** | |
| [%] | 0 | 100 |

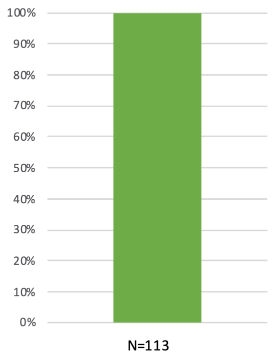

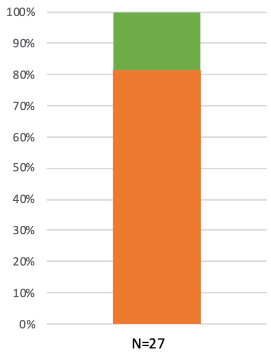

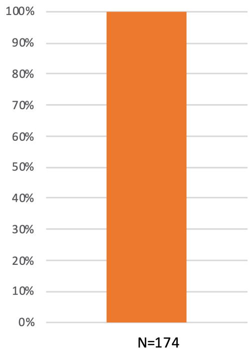

**Supplementary Fig. 6:**

Source data underlying the graphs in Fig. 3b showing the distribution of either success or failure of FtsZ ring/divisome progression among the cells counted. n.e., not evaluated.

### Supplementary Table 1

Source data underlying the graphs of *in vitro* FtsZ polymerization according to Fig. 1b.

| | no GTP, no ADEP | | | plus GTP, no ADEP | | | plus GTP, plus ADEP | | |
|---|---|---|---|---|---|---|---|---|---|
| time (min) | replicate 1 (transmission %) | replicate 2 (transmission %) | mean (transmission %) | replicate 1 (transmission %) | replicate 2 (transmission %) | mean (transmission %) | replicate 1 (transmission %) | replicate 2 (transmission %) | mean (transmission %) |
| -4 | 100.0 | 100.0 | 100.0 | 100.0 | 100.0 | 100.0 | 100.0 | 100.0 | 100.0 |
| -2 | 102.3 | 99.2 | 100.8 | 102.3 | 99.2 | 100.8 | 102.3 | 99.2 | 100.8 |
| 0 | 101.7 | 99.2 | 100.5 | 101.7* | 98.1* | 99.9 | 101.7* | 98.1* | 99.9 |
| 2 | 102.2 | 99.2 | 100.7 | 100.6 | 82.9 | 91.8 | 100.6 | 82.9 | 91.8 |
| 4 | 101.0 | 99.5 | 100.3 | 86.1 | 68 | 77.1 | 86.1 | 68 | 77.1 |
| 6 | 101.4 | 99.4 | 100.4 | 71.5 | 59.4 | 65.5 | 71.5 | 59.4 | 65.5 |
| 8 | 102.3 | 99.7 | 101.0 | 59.9 | 55.8 | 57.9 | 59.9 | 55.8 | 57.9 |
| 10 | 102.1 | 99.1 | 100.6 | 46.4 | 54.3 | 50.4 | 46.4 | 54.3 | 50.4 |
| 12 | 102.2 | 99.0 | 100.6 | 51.4 | 54.3 | 52.9 | 51.4 | 54.3 | 52.9 |
| 14 | 102.7 | 99.0 | 100.9 | 49.8 | 54.5 | 52.2 | 49.8 | 54.5 | 52.2 |
| 16 | 101.6 | 97.7 | 99.7 | 48.2 | 54.9 | 51.6 | 48.2 | 54.9 | 51.6 |
| 18 | 102.8 | 99.1 | 101.0 | 51 | 55.6 | 53.3 | 50.4** | 52.8** | 51.6 |
| 20 | 103.3 | 99.4 | 101.4 | 49.4 | 55.4 | 52.4 | 49.2 | 52.9 | 51.1 |
| 25 | 100.8 | 99.4 | 100.1 | 47.7 | 55.1 | 51.4 | 48 | 52.6 | 50.3 |
| 30 | 100.7 | 98.0 | 99.4 | 49.3 | 54.1 | 51.7 | 47.1 | 54.2 | 50.7 |
| 35 | 100.9 | 99.2 | 100.1 | 48.7 | 55.9 | 52.3 | 48.4 | 55.7 | 52.1 |
| 40 | 100.3 | 99.3 | 99.8 | 51.8 | 57.8 | 54.8 | 49.9 | 54.3 | 52.1 |
| 50 | 101.3 | 99.9 | 100.6 | 52.2 | 56.9 | 54.6 | 48.4 | 53.9 | 51.2 |

*addition of GTP, **addition of ADEP
